# Fast Detection of Differential Chromatin Domains with SCIDDO

**DOI:** 10.1101/441766

**Authors:** Peter Ebert, Marcel H. Schulz

## Abstract

The generation of genome-wide maps of histone modifications using chromatin immunoprecipitation sequencing (ChIP-seq) is a common approach to dissect the complexity of the epigenome. However, interpretation and differential analysis of histone ChIP-seq datasets remains challenging due to the genomic co-occurrence of several marks and their difference in genomic spread. Here we present SCIDDO, a fast statistical method for the detection of differential chromatin domains (DCDs) from chromatin state maps. DCD detection simplifies relevant tasks such as the characterization of chromatin changes in differentially expressed genes or the examination of chromatin dynamics at regulatory elements. SCIDDO is available at github.com/ptrebert/sciddo

## Background

Large epigenome mapping consortia such as DEEP [1], BLUEPRINT [2] or ENCODE [3] produce an ever–increasing amount of reference epigenomes for a multitude of different cell types. With the ultimate goal of compiling a publicly available catalog of 1000 reference epigenomes released under the IHEC [4] umbrella, the computational interpretation of large amounts of epigenome data presents a formidable challenge for bioinformatics. However, the cell–type specific and dynamic nature of the epigenome adds substantial complexity to the problem of characterizing cellular similarities and differences on the epigenetic level. Moreover, limited resources commonly force scientists to investigate only a small number of biological replicates per condition of interest. Despite all these challenges, the discoveries in the field of epigenomics have greatly enhanced our understanding of transcriptional regulation, cellular identity and disease development [5–9].

An important component of the epigenetic landscape are post–translational modifications of histone proteins, briefly referred to as histone marks. The interpretation of histone mark data is particularly intricate as the interplay between different histone marks results in a combinatoric complexity that is largely absent for other epigenetic modifications such as DNA methylation. To give an example, bivalent chromatin domains that mark developmental genes in embryonic stem cells represent a biologically meaningful co–occurrence of several different histone marks [10, 11]. The realization that histone mark combinations can be interpreted as local activity states of the genome, so–called chromatin states, led to the widespread use of probabilistic graphical models to discover these “hidden states” [12–16]. Popular tools such as ChromHMM [13] or EpiCSeg [15] have tremendously simplified the analysis of histone data as they summarize the combined effect of histone mark co–occurrences in a manageable number of discrete chromatin states. After functional characterization, the discovered chromatin states are commonly augmented with textual labels to ease interpretation, e.g., identifying regions as active or poised promoters, or distinguishing between weak and strong transcriptional activity. However, in our experience, the generated chromatin state maps are often manually inspected in only a limited number of loci or simply serve as additional genomic annotation data. Given that chromatin state maps provide a neat abstraction of the various histone mark combinations, it stands to reason that a more comprehensive view on them may offer valuable guidance in exploratory studies.

So far, there are only few tools available that use chromatin state maps to identify regions of differential chromatin marking. ChromDet [17] can be applied in a genome–wide manner and uses multiple correspondence analysis (an analog to principal component analysis for categorical data) followed by an iterative clustering approach to identify regions that perfectly partition the samples into cell–type or lineage specific groups (so–called chromatin determinant regions). The computational burden of a ChromDet analysis is lowered by various filtering steps to remove uninformative or outlier regions, which renders ChromDet analyses prohibitive for small sample numbers. This preprocessing also requires enough insight into the nature of the samples at hand to manually set appropriate filtering thresholds.

Other available tools for the differential analysis of chromatin state maps enable only the analysis of a predefined set of genomic regions. The ChromDiff [18] tool represents chromatin states in user–specified regions of interest, e.g., the bodies of all coding genes, as normalized coverage vectors. After batch effect correction using a regression model, ChromDiff uses the non–parametric Mann–Whitney–Wilcoxon test to identify differential chromatin states between sample groups, e.g., contrasting all male and female samples. Since ChromDiff relies on standard statistical tests for its analysis, sufficient statistical power in terms of number of available samples per group is mandatory to find any significant differences between the groups. The recently published Chromswitch package [19] similarly identifies differential chromatin states only in preselected regions of interest. Chromswitch can only analyze a single chromatin state at a time and uses a binary “presence/absence” encoding to construct feature vectors that are subsequently clustered. The cluster assignments resulting from the hierarchical clustering are then scored by their agreement with the known biological labels of the samples and manual thresholding on these score is required to select the final set of chromatin state switches.

A common denominator of all existing methods is that they consider chromatin state similarity as a binary variable, i.e., any chromatin state is, to exactly the same extent, (dis–) similar to any other chromatin state. We argue that this is an oversimplification and, as we will show below, much is to be gained when measuring chromatin state similarity using a quantitative scale.

In summary, current methods are limited to region–based analysis, focus on individual chromatin states, require a comparatively large number of biological replicates for their statistical analysis, and use a quite basic representation of chromatin state similarity, which hinders general applicability of existing methods.

We devised a new method for the score–based identification of differential chromatin domains (SCIDDO) with the goal of providing a generally applicable tool for the fast identification of differential chromatin marking. One of SCIDDO’s main features is its capability to identify potentially large and heterogeneous regions of differential chromatin marking, which we refer to as differential chromatin domains (DCDs). The statistical evaluation of the identified domains relies on well–established theory borrowed from score–based biological sequence analysis. This transfer of theory enables an interpretable presentation of SCIDDO’s results and facilitates downstream analysis. In the following, we present results obtained by analyzing four groups of replicated human samples with SCIDDO. In this straightforward analysis, we assessed the robustness of our method by comparing DCDs between individual replicates and characterized the identified domains by overlapping them with differentially expressed genes (DEGs) and various regulatory annotation datasets. We compared SCIDDO to other methods for the differential analysis of histone data and collected evidence that highlights SCIDDO’s usefulness in identifying regions of dynamic chromatin changes, e.g., enhancers switching from an “on” to an “off” state between cell types. Finally, we discuss potential limitations and future applications of our method.

## Results

### Score–based identification of differential chromatin domains

The differential analysis with SCIDDO consists of two major parts, data preparation and the actual analysis run (see Figure 1 for an overview). In the data preparation step (Figure 1 (A)), SCIDDO creates a single coherent dataset storing all data and metadata relevant for the analysis to ensure later reproducibility of the results. As part of the data preparation, the state emission probabilities of the chromatin state segmentation model are used to compute pairwise chromatin state dissimilarities (see Methods). Starting from this dataset, SCIDDO then performs the differential analysis as follows: for each comparison contrasting sample group X versus group Y, SCIDDO first compares individual replicates against each other, say, X–2 versus Y–1 (Figure 1 step (B)). In this process, each observed chromatin state pair in the two chromatin state maps is assigned a score that quantifies the dissimilarity of the two states: positive scores indicate state dissimilarity, and negative scores indicate state similarity (Figure 1 (C); see Methods). Candidate regions showing differential chromatin marking are identified on this level of replicate comparisons by searching for chromosomal segments that show a high cumulative score, which indicates a strong dissimilarity on the chromatin state level; hence, we refer to this value as the differential chromatin score (DCS) of the segment (Figure 1 step (C) to (D)). It should be pointed out that extracting segments based on (locally) maximal DCSs implies also a maximization of the segment length, and no (predefined) minimum or maximum length has to be specified. To proceed from candidate regions identified in individual replicate comparisons (e.g., X–2 versus Y–1) to candidate regions that are representative of all samples X versus Y, overlapping candidate regions are merged by averaging their DCSs and taking the union of their genomic coverages (Figure 1 (E)). As the final step in the analysis, the segment DCSs are turned into an Expect (E) value, which allows to filter the resulting candidate regions for their statistical significance (Figure 1 step (F)). The E–value (see Methods) has the interpretation of indicating how many candidate regions with at least a similarly high DCS could arise simply due to chance when comparing random sequences of the same length. In other words, when filtering the candidate regions for a default E–value of less than 1 to call DCDs, SCIDDO restricts the results to those chromosomal regions where the chromatin states are so different between the samples that one would not expect to find such a difference simply due to chance. To simplify visualizations, we report E–values after a negative log10 transform in the remainder of this study. The aforementioned threshold of 1 is thus transformed to 0 and larger E–values indicate higher statistical stringency.

**Figure 1:**
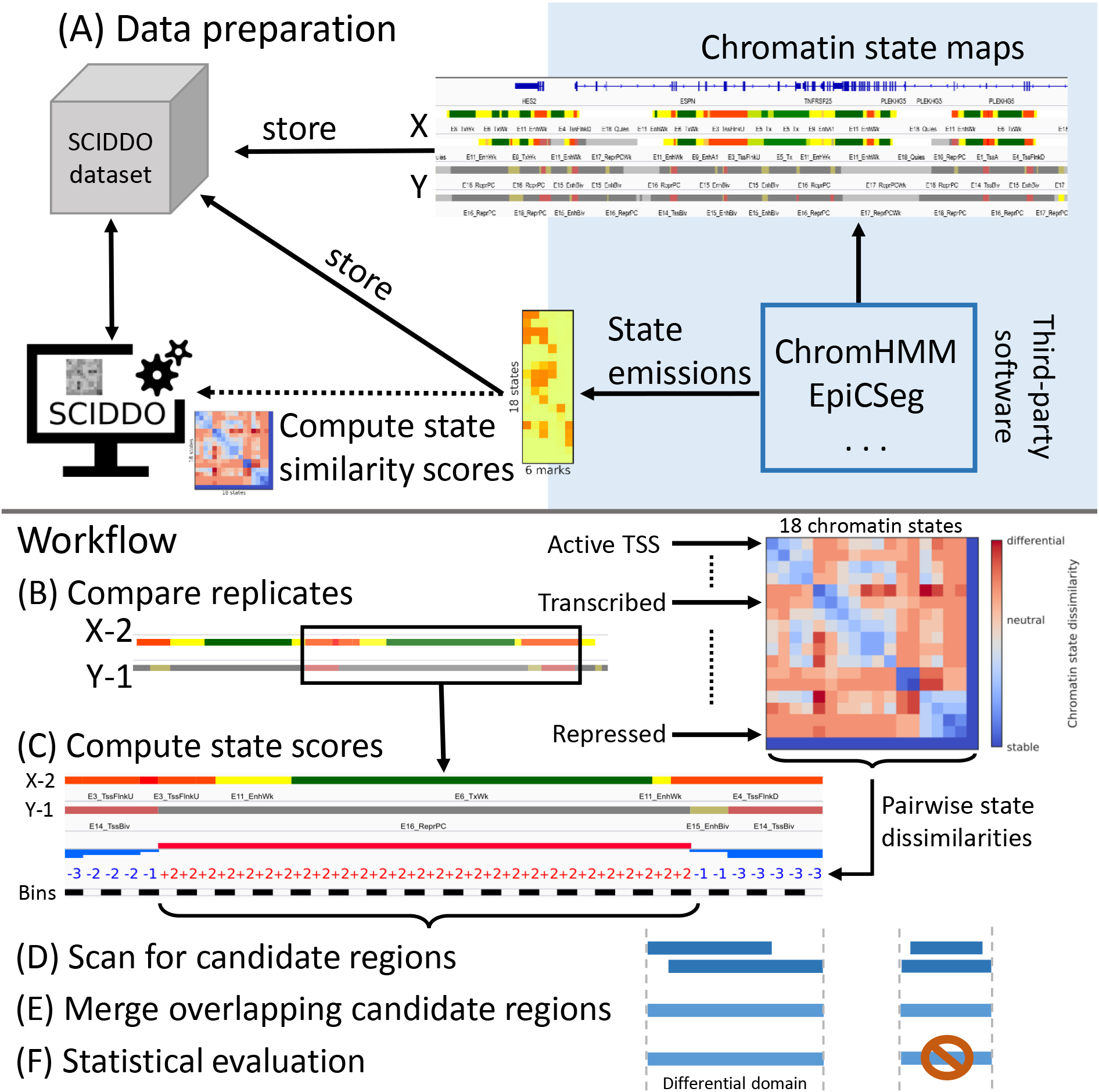
Overview of SCIDDO’s workflow to identify differential chromatin domains. (A) Data preparation: chromatin state maps can be generated using common tools (blue shaded area). The chromatin state maps for all replicates of sample groups X and Y are stored together with the chromatin state emission probabilities in a SCIDDO dataset to ensure later reproducibility of the analysis. The state emission probabilities are used to compute chromatin state similarity scores. (B)–(F) Workflow: the differential analysis starts by comparing all replicate pairs in the dataset, here exemplified as X–2 vs. Y–1 (B). All observed chromatin state pairs are scored with their respective dissimilarity score (C). The resulting score sequences are scanned for high–scoring candidate regions (D). Overlapping candidate regions of all replicate pairs are then merged (E) and filtered after statistical evaluation to generate the final set of differential chromatin domains
(F).

To demonstrate the usefulness of SCIDDO in a differential analysis setting, we compiled a medium–sized dataset of high quality DEEP samples that includes both distantly as well as more closely related cell types. We selected three biological replicates of monocytes (Mo 1, 3 and 5) and two biological replicates of macrophages (Ma 3 and 5); these hematopoietic samples have been extensively characterized in previous work [20] and are only separated by a single step of cellular differentiation. Additionally, we selected two biological replicates of hepatocytes (He 2 and 3; Additional file 1: Table S1) and two replicates of the HepG2 cell line (HG 1 and 2; Additional file 1: Table S1). Though HepG2 is commonly used as an *in vitro* model in liver–related studies, its state as an immortalized cell line distinguishes it from the primary hepatocytes in our dataset. Hence, the dataset we compiled enabled us to evaluate SCIDDO’s performance at various degrees of “cellular relatedness”.

For each sample in the dataset, we included the six histone marks (H3K27ac, H3K27me3, H3K36me3, H3K4me1, H3K4me3, H3K9me3) plus the respective Input control that are defined as the chromatin constituents of an IHEC reference epigenome (Additional file 1: Table S1). For later functional characterization of the identified DCDs, we also downloaded mRNA–seq expression data for all samples (Additional file 1: Table S2). The chromatin state maps that we used for the following analysis were generated using a predefined ChromHMM model (CMM18) [12, 13, 21] to simplify interpretation of the chromatin states (see Methods; Additional file 1: Figure S1, Table S3). Scores representing pairwise dissimilarity between chromatin states were derived from the chromatin state emission probabilities of the same ChromHMM model (see Methods).

We performed a differential analysis for all six possible pairings of sample groups in our dataset, i.e., (i) HepG2 vs. hepatocytes; (ii) HepG2 vs. monocytes; (iii) HepG2 vs. macrophages; (iv) hepatocytes vs. monocytes; (v) hepatocytes vs. macrophages, and (vi) monocytes vs. macrophages. The entire SCIDDO analysis including data preparation completed within minutes on a moderately powerful compute server (Additional file 1: Table S4). The results presented for this analysis are structured as follows: first, we provide some evidence that our data follow the theoretical assumptions necessary for a sound statistical evaluation. Next, we highlight the robustness of SCIDDO’s results across replicates and then provide a more biology–oriented characterization of the identified DCDs.

### Differential chromatin scores follow extreme value distribution

The last step in the SCIDDO workflow described above consists of turning the DCSs into an E–value that is used for filtering the set of candidate regions to obtain the final set of DCDs. This step relies on theory developed for biological sequence analysis (see Methods) and requires first a normalization of the raw cumulative DCSs to account for the fact that comparing longer chromosomal sequences increases the chances of observing higher cumulative DCSs. This normalization uses two estimated statistical parameters, λ and *K*, that lack a biological interpretation, but can be thought of as scaling factors for the scoring system and the sequence length, respectively. Second, the theory assumes a null model of random sequences, and under this null model, the distribution of the scores should in the limit converge in distribution to a Gumbel–type extreme value distribution (see Methods). We confirmed that this is indeed the case in our analysis by comparing randomized chromatin state maps with each other and fitting all maximal DCSs identified during this sampling procedure to a Gumbel distribution (Figure 2A). We also plotted the per–chromosome estimates of the statistical parameters λ and *K* that are needed for the score normalization (Figure 2B; see Methods), and could confirm that the estimates are within reasonable bounds given examples from literature [22]. The observed agreement with theory thus supports the last step in the SCIDDO analysis (Figure 1 step (C)) of filtering candidate regions based on their E–value.

**Figure 2:**
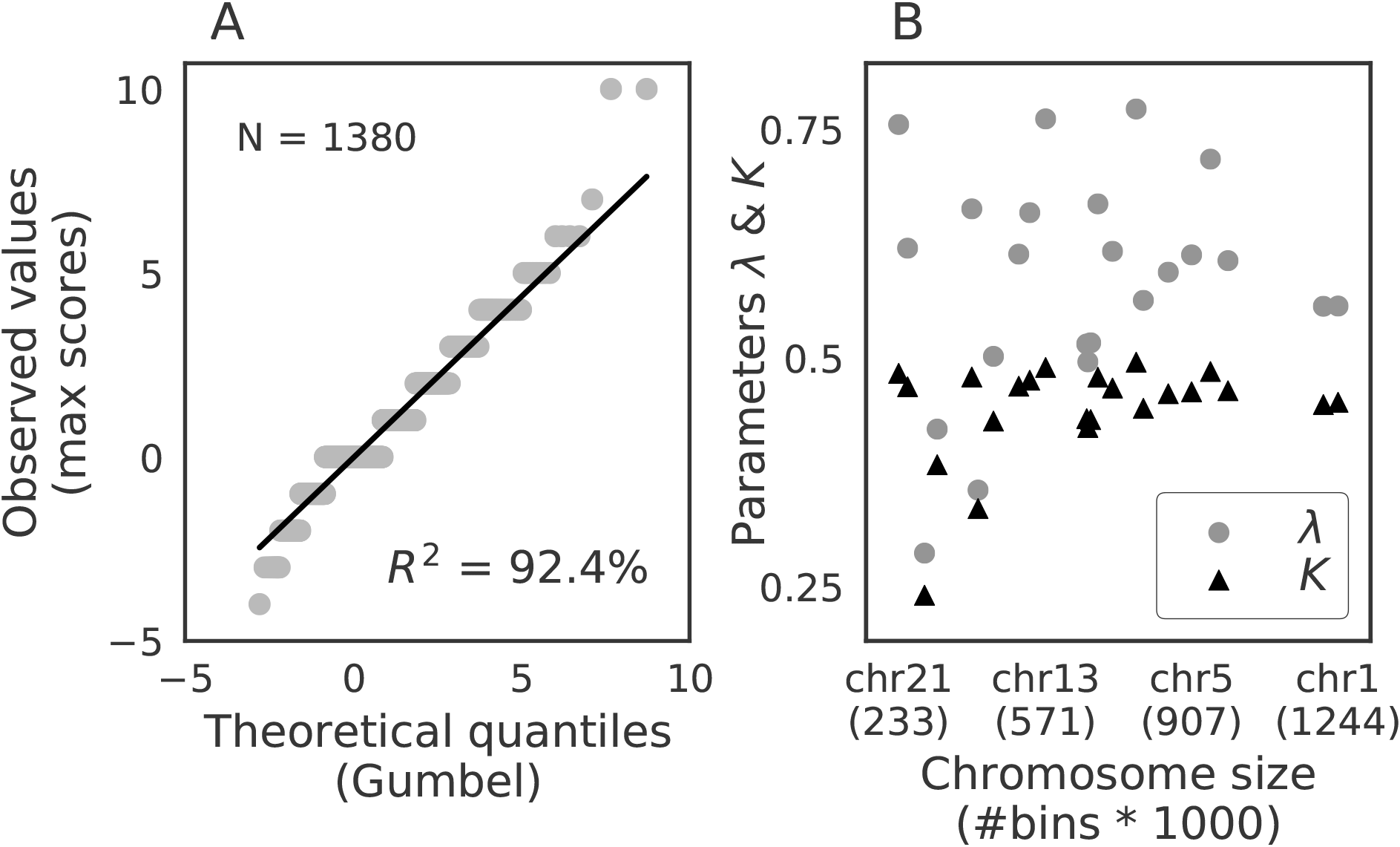
Observed maximal scores and parameter estimates follow theoretical assumptions. (A) Probability plot of all normalized maximal scores derived from comparing random sequences (y–axis) fit to the theoretical quantiles of a Gumbel–type extreme value distribution (x–axis). (B) Chromosomes are sorted by increasing size (in genomic bins) from left to right (x–axis) and the per–chromosome estimates of the two statistical parameters λ (gray points) and *K* (black triangles) are plotted on the same scale (y–axis). *R*^2^: coefficient of determination

### SCIDDO robustly identifies differential chromatin domains

Histone ChIP–seq data is known to be affected by various sources of noise, e.g., ranging from artifacts introduced during library preparation, to irregularities caused by varying mappability in the reference genome or to spurious signal due to unspecific antibody binding [23–25]. In combination, biological and technical variation can render any differential analysis pointless if the results are dominated by noise, and not by the biological signal of interest. To test if the identified candidate regions were indeed representative and not replicate–specific, we computed the Spearman correlation of the E–values between all overlapping candidate regions. We visualized an exemplary case selected based on the mean of all comparisons. This exemplary case shows a Spearman correlation of 0.72 between the candidate regions (Figure 3). The red bars in the lower left corner indicate candidate regions that are unique to the respective replicate comparison. It can be observed that unique candidate regions tend to have comparatively lower E–values whereas those candidate regions found in both replicate comparisons tend to have higher E–values. In general, the average Spearman correlations across all replicate comparisons are consistently in high range from 0.67 (HepG2 vs. hepatocytes) to 0.73 (HepG2 vs. monocytes; Additional file 1: Table S5).

**Figure 3:**
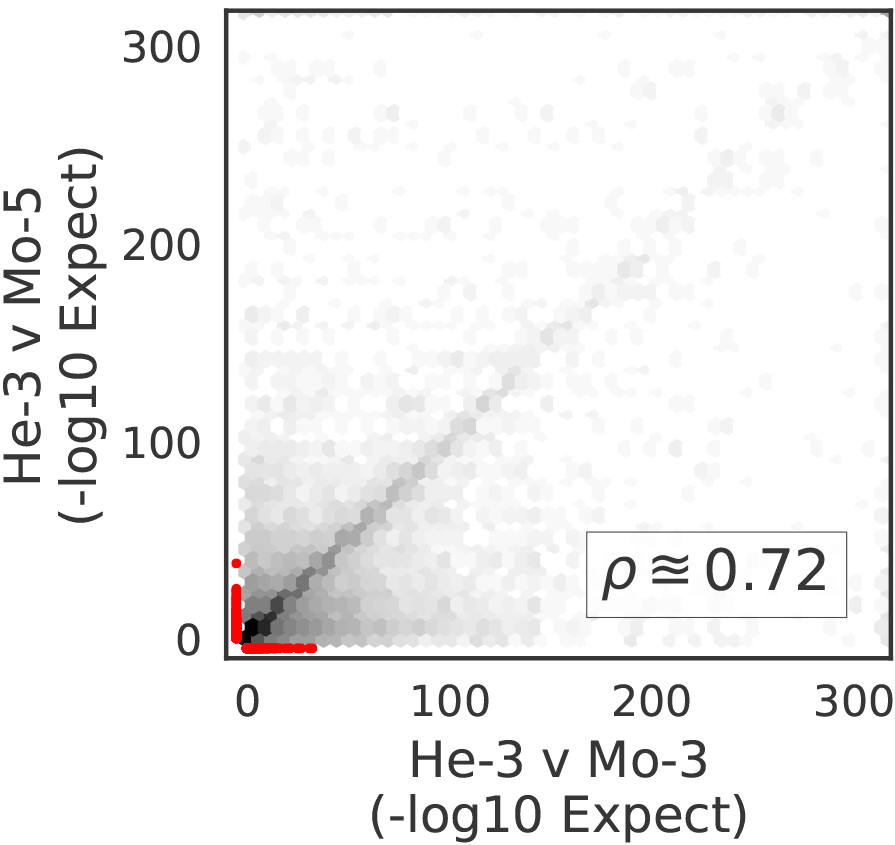
Candidate regions are robustly identified across individual replicates. Exemplified agreement of candidate regions identified in replicate comparisons. E–values of candidate regions identified for He–3 vs. Mo–3 (x–axis) are plotted against E–values of overlapping candidate regions identified for He–3 vs. Mo–5 (y–axis). The red area indicates E–values of those candidate regions that are unique to the respective replicate comparison. *ρ*: Spearman correlation of E–values

### Differential chromatin domains occur in various regulatory contexts

Since it is well–established that histone marks occur in various regulatory contexts, e.g., ranging from promoters and enhancers to gene bodies, it stands to reason that *bona fide* DCDs should predominantly occur in similar regulatory contexts. To test this hypothesis, we intersected the DCDs identified by SCIDDO with various annotation datasets and observed that, in general, around 80 to 90% of all DCDs overlap with at least one type of genomic annotation (Figure 4). Since there is no theory that would enable us to formulate an *a priori* expectation about the extent to which differences on the chromatin level should occur between any two cell types, we cannot assess the plausibility of the absolute numbers of identified domains. Nevertheless, it can be observed that the lowest number of domains is detected in the comparison of monocytes to macrophages (Figure 4F), i.e., when comparing the two most closely related cell types in our dataset. For all other comparisons, the number of identified chromatin domains is approximately 4–to more than 5–fold higher, but yet shows a similar tendency of a smaller number of identified chromatin domains for more closely related cell types.

**Figure 4:**
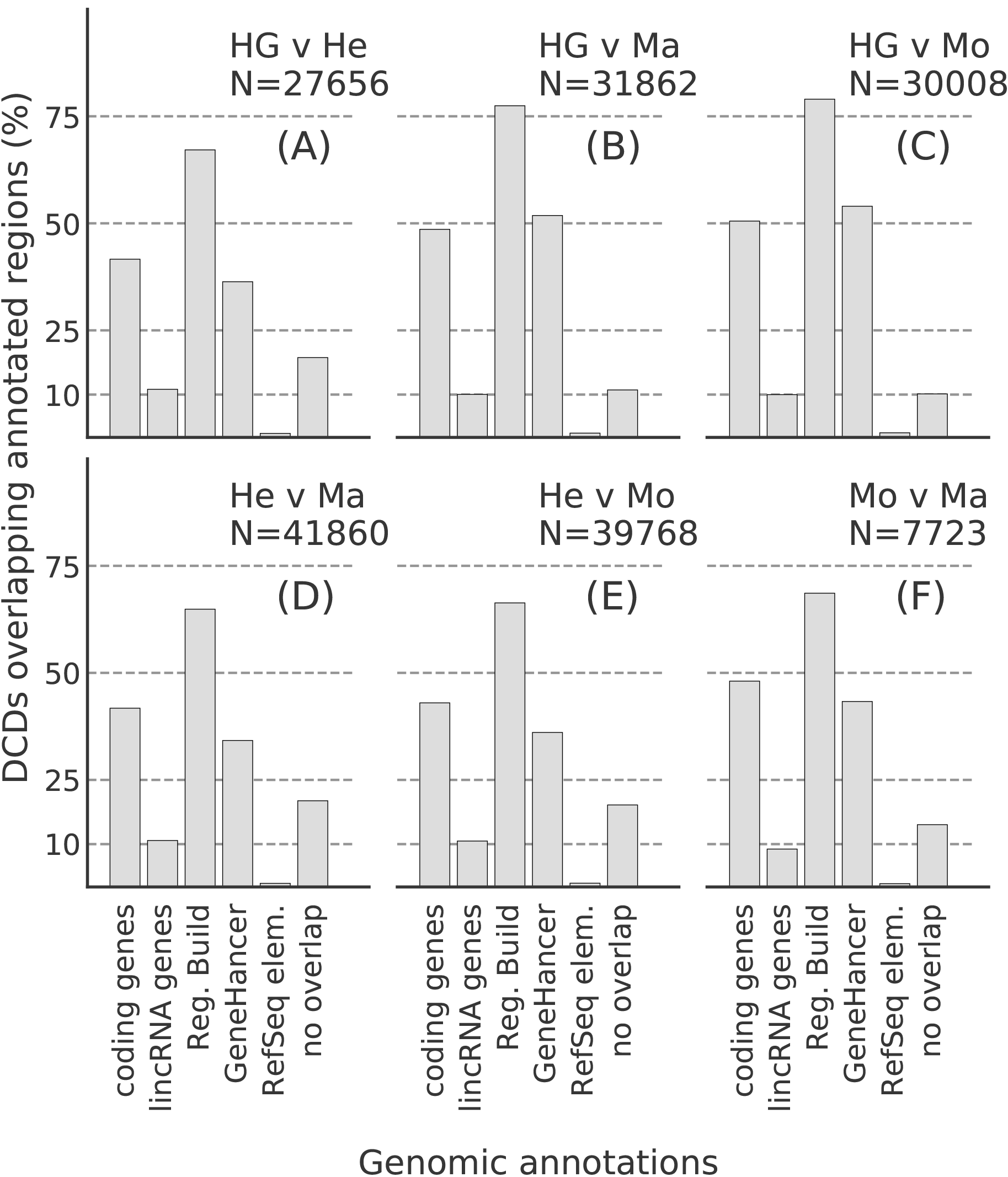
Differential chromatin domains overlap with annotated regulatory regions. Bar heights indicate percentage of identified differential chromatin domains that overlap with different genomic annotations for all six sample group comparisons (A–F). *N*: total number of identified domains; coding genes: Gencode v21 protein–coding genes; lincRNA genes: Gencode v21 lincRNA genes; Reg. Build: Ensembl Regulatory Build v78; GeneHancer: GeneHancer annotated enhancers limited to Gencode v21 gene set; Refseq elem.: RefSeq functional elements

These results also illustrate that the distribution of overlaps seems not to be affected by the number of DCDs identified. In all comparisons, at least ~70% of the DCDs overlap with at least one regulatory region annotated in the Ensembl Regulatory Build [26]. The Regulatory Build comprises several different types of regulatory regions and has extensive genome coverage. Hence, the Regulatory Build enables us to interpret the relevance of DCDs in light of various functional categories. Since the distribution of genomic locations of the DCDs seems fairly similar across all comparisons, and analogous observations can be made when examining the length distribution of the DCDs (Additional file 1: Figure S2), we examined if there is a difference in DCD E–values aggregated over all comparisons (Figure 5). DCDs overlapping any regulatory region show higher E–values compared to those DCDs that have no overlaps (Figure 5, bottom panel). This effect is most pronounced for annotated promoters and transcription factor binding sites (TFBS), and this seems not to be an effect of regulatory region size (Figure 5, top panel). The average number of distinct regulatory region overlaps per DCD shows that a DCD often spans several of the shorter regulatory regions, with the exception of TFBS, which is the least abundant region type with a median size <1 kbp in the Regulatory Build. At the other end of the size spectrum are promoters, which also show hardly any variation around a median of one DCD overlap per promoter.

**Figure 5:**
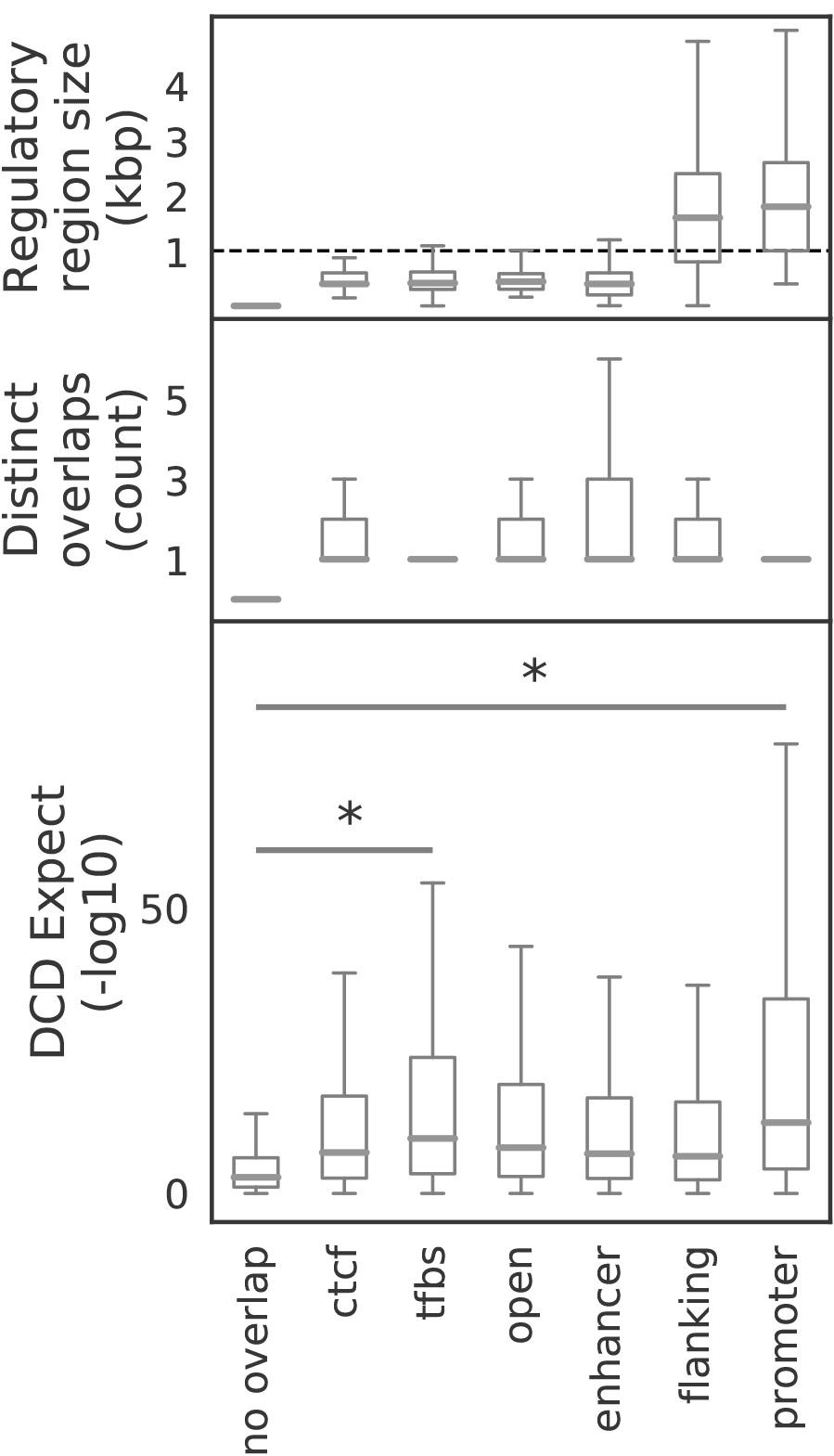
E–value distribution of DCDs overlapping regulatory regions. Bottom: boxplots show distribution of E–values of all differential chromatin domains overlapping regulatory region types as annotated in the Ensembl Regulatory Build (v78) aggregated over all sample comparisons. Differences in magnitude of E–values were assessed with two–sided Mann–Whitney–U test and considered significant (*) at *p* < 0.01. Middle: boxplots show distinct overlaps per DCDs, i.e., the number of regulatory regions of that type overlapping the same DCD. Top: Size distribution of the Ensembl regulatory regions. Dashed line indicates a size of 1000 bp. Regulatory region types: ctcf: CTCF binding sites; tfbs: transcription factor binding sites; open: regions of open chromatin; enhancer: enhancer; flanking: promoter–flanking regions; promoter: promoter

### Formation of differential chromatin domains affects gene expression

The results presented in the previous section indicate that DCDs largely overlap with a variety of regulatory regions, and thus it seems plausible that the formation of a DCD should have functional consequences, e.g., by modulating gene expression levels. Apart from basic considerations about the magnitude of the observed E–values, we also hypothesized that DCDs covering larger parts of the gene body could indicate stronger changes in gene expression. To give a canonical example, a gene that is entirely repressed by means of polycomb–mediated silencing should be enriched for the histone mark H3K27me3, and this marking should be replaced by H3K36me3 as soon as the gene is activated and actively transcribed [27]. On the other hand, if the gene expression is modulated, e.g., by changing transcription factor binding in enhancer regions, the effect on the chromatin marking in the gene body could arguably be less pronounced. To investigate this hypothesis, we stratified all genes by the fraction of their gene body length being covered by a DCD (no overlap in gene body or enhancers, less or more than 50% gene body overlap). Next, we computed gene expression fold changes using DESeq2 [28] (see Methods) for the six sample group comparisons and visualized the fold change for all genes in the three DCD overlap groups as a cumulative distribution (Figure 6; Additional file 1: Figures S3 and S4). The curves indicate that genes covered by more than 50% of their body length with a DCD indeed exhibit stronger changes in their expression level (orange lines). A similar effect, albeit weaker, can be observed for genes having less than 50% of their body or their promoter covered by a DCD (blue lines). In many cases, the difference in fold change relative to the group of genes that does not overlap a DCD is significant. Additionally, we applied the same method to test if the number of gene–associated enhancers that overlap a DCD had a similar bearing on gene expression (Figure 6, middle and right panels; Additional file 1: Figures S3 and S4). This enhancer–centric view shows a stable pattern across most sample comparisons that indicates that stronger changes in gene expression occur if more gene–associated enhancers overlap a DCD. This observation is particularly intriguing when restricting the view on intergenic enhancers, where, as opposed to intragenic enhancers, there is lower chance of a coincidental overlap with a DCD. In general, a small but noticeable difference compared to the no DCD overlap group (gray dashed line) can be expected as soon as 2–3 enhancers show a DCD (magenta curve).

**Figure 6:**
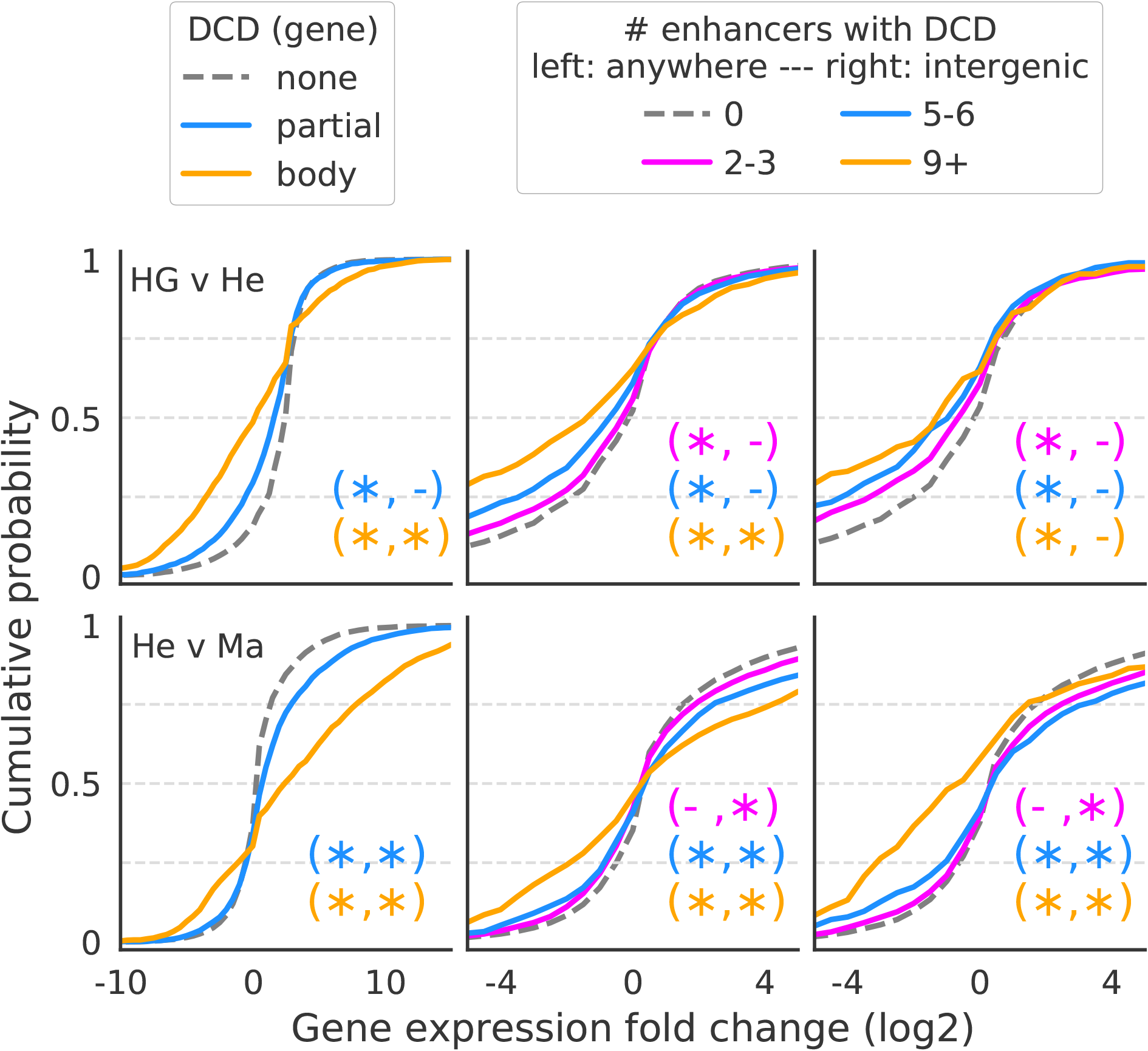
DCDs overlapping gene bodies and enhancers affect gene expression. Left panels: genes were stratified by the amount of DCD overlap either covering more than 50% of the body (body; orange curve) or less than 50% of the body or the promoter region (partial; blue curve). Expression fold change of the genes in the respective groups is plotted along the x–axis within a restricted window for improved readability. Statistical significance of the difference in mean fold change of the groups relative to the no overlap group (“none”) was computed separately for negative and positive fold change genes using a two–sided Mann–Whitney–U test (“*” significant with *p* < 0.01, “−” not significant otherwise). Middle and right panels: same analysis as for the gene body, but here counting the number of intra– and intergenic enhancers (anywhere, middle) or only intergenic enhancers (right) per gene that overlap a DCD. Expression fold changes plotted within a restricted window for improved readability. Statistical significance assessed as above.

### SCIDDO detects chromatin changes in differentially expressed genes

By design, SCIDDO does not impose any restrictions on the regions of interest that can be interrogated in a differential analysis. Since there is no general model of chromatin variation that would enable us to assess the plausibility of the identified differential chromatin domains irrespective of their genomic context, we decided to focus on a small–scale case study that is arguably of broad biological interest.

We investigated to what extent DCDs can be used to specifically identify differentially expressed genes (DEGs). As ground truth for this analysis, we used the same DESeq2 results as above, but applied a threshold to split the genes into differentially expressed and stable ones (see Methods). As a first step, we checked what percentage of DEGs could be recovered using SCIDDO’s DCDs (Figure 7). For four out of the six sample comparisons, more than 90% of all DEGs could be recovered with DCDs either overlapping the gene body, the gene promoter or at least one gene–associated enhancer. For the comparison of HepG2 to primary hepatocytes (Figure 7A), approximately 81% of DEGs could be recovered, and for the comparison of monocytes to macrophages, 54% of all DEGs were recoverable by using DCDs (Figure 7F). The comparatively lower rate of DEG recovery for the monocyte to macrophage comparison seems to be in line with the already observed trend of fewer differences on the chromatin level with increasing cellular relatedness (e.g, see Figure 4). We present a more in–depth analysis of this observation in the following section of the Results. Next, we tested if it was possible to broadly distinguish between DEGs and stably expressed genes by thresholding on the E–values of the DCDs that overlap gene bodies. To that end, we stratified the set of DEGs based on their fold change into three groups (top 20%, middle and bottom 40%) and plotted the E–value distribution of the DCDs for these three groups and for all other chromatin domains (Figure 8, bottom panel). We find that DEGs with the highest fold change in expression overlap DCDs that have a significantly higher E–value on average relative to DCDs overlapping the remaining DEGs. Furthermore, it is interesting to observe that the E–value distribution of the DCDs overlapping stable genes is similar to those that do not overlap any gene (but could, e.g., overlap with intergenic enhancers). The number of distinct DCDs that overlap any given gene shows no notable variation across all groups (Figure 8, middle panel). The distribution of the gene body lengths in the respective groups appears to be fairly balanced (Figure 8, top panel) and thus does not suggest that the number of DCD overlaps or the observed difference in E–value distribution is a simple effect of gene body length. We explicitly confirmed this by repeating the analysis, but this time stratifying DEGs by gene body length (Additional file 1: Figure S5). The E–values of the DCDs overlapping the longest genes are comparatively lower, and this suggests that larger E–values are probably not a result of increasing gene body length.

**Figure 7:**
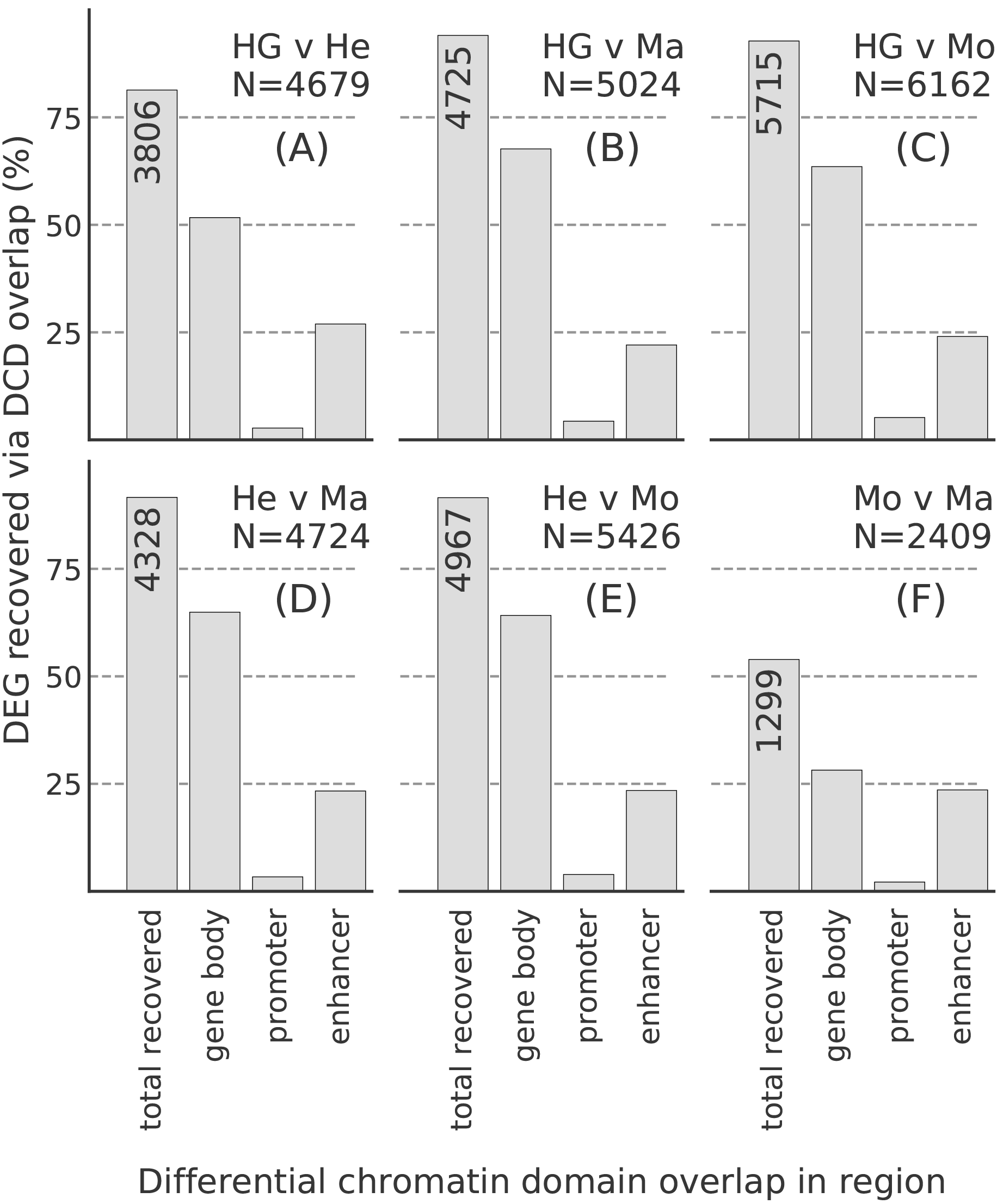
Differential chromatin domains recover differentially expressed genes. Bar heights indicate percentage of recovered differentially expressed genes by counting overlaps with differential chromatin domains in gene bodies, in gene promoters (but not in gene bodies) or in gene–associated enhancers (but not in gene bodies or gene promoters). The leftmost bar is annotated with the total number of recovered genes. *N*: total number of differentially expressed genes per comparison A–F.

**Figure 8:**
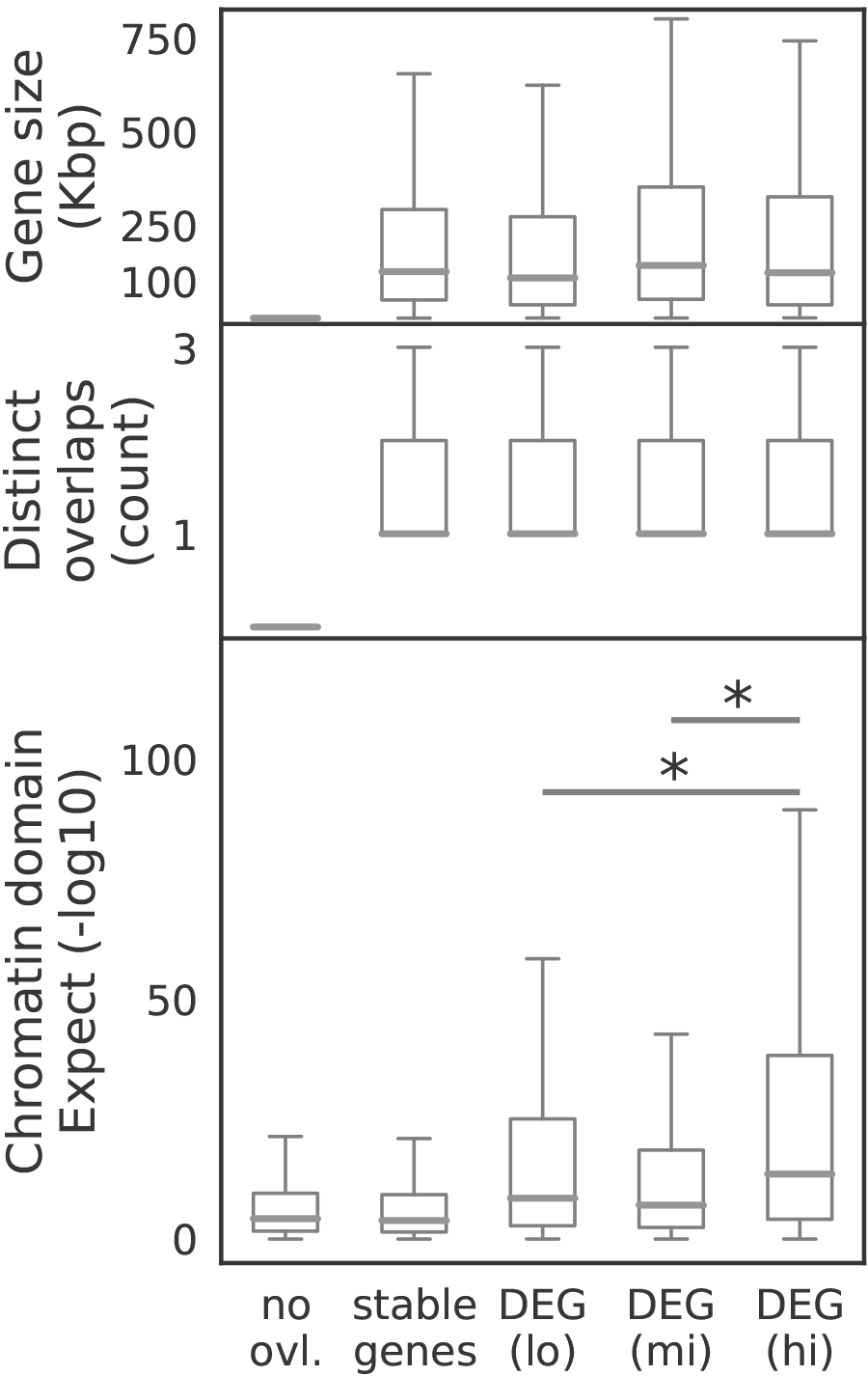
E–value distribution of DCDs overlapping gene bodies. Genes were stratified into four groups based on their expression fold change (stable/no change, lowest 40%, middle 40% and top 20% of DEGs according to expression fold change). Bottom: boxplots show distribution of E–values of all DCDs overlapping gene bodies in the respective groups aggregated over all sample comparisons. The no overlap group contains all E–values of DCDs not overlapping any gene. Middle: boxplots show distinct DCD overlaps per gene. Top: boxplots show gene body length distribution of all genes in the respective group. Differences in magnitude of E–values were assessed with a two–sided Mann–Whitney–U test and considered significant (*) with *p* < 0.01.

### Methodological and biological limitations for chromatin–based detection of differentially expressed genes

The theory borrowed from local scoring and implemented in SCIDDO is used to assign a measure of statistical stringency — the E–value — to each discovered DCD. Yet, the theory does not offer a way to decide what threshold on the E–value best separates genuine from chance observations. The necessary normalization to account for the length of the sequences being compared immediately suggests that short but biologically *true* differential regions will be assigned an (untransformed) E–value well above SCIDDO’s default threshold of 1.

We checked the extent to which the default E–value threshold of 1 could limit SCIDDO’s ability to identify — especially short — DEGs. We binned all DEGs by their gene body size and plotted the amount of genes with a DCD overlapping their gene body at E–value thresholds of 1 and 100 (Figure 9). The histogram shows the expected behavior of SCIDDO to predominantly recover longer DEGs by means of finding a DCD in their gene body. However, relaxing the E–value threshold seems not to affect this general trend as the additional DEGs also show a tendency toward longer gene bodies. We thus wondered if other technical or biological artifacts might exacerbate the detection of DEGs on the chromatin level. We focused specifically on the comparison of monocytes to macrophages where approximately only 54% of all DEGs could be recovered using DCDs (see Figure 7F).

**Figure 9:**
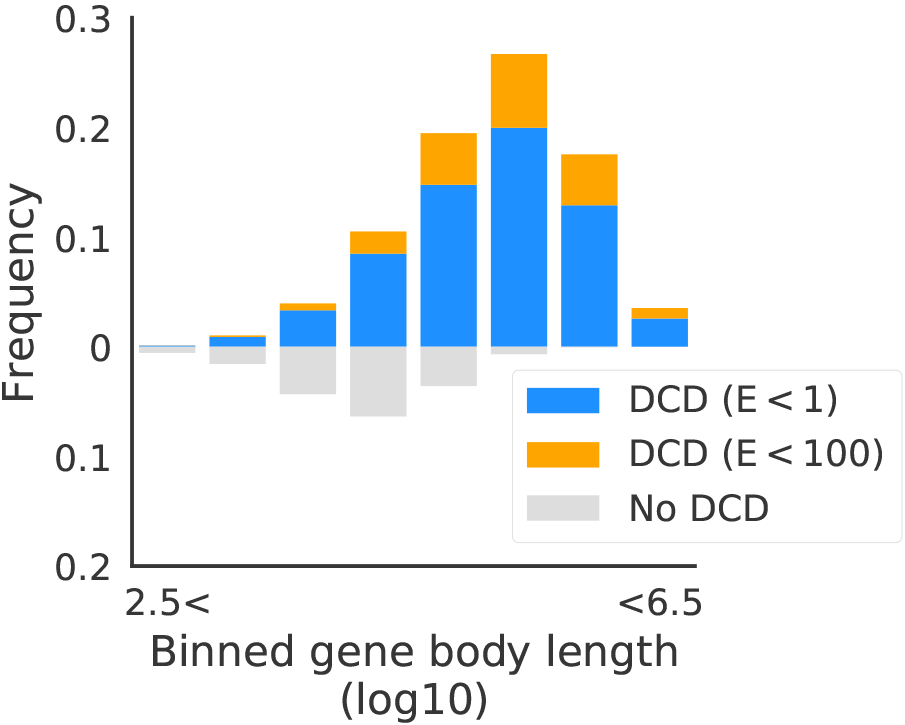
Relaxing E–value threshold does not help in detecting short DEGs. All DEGs for all six comparisons were binned based on their gene body size (x–axis) and classified based on overlapping DCDs in their gene body (y–axis). DCDs were called with the default threshold of E < 1 (blue) and with a relaxed threshold of E < 100 (orange).

As a first step, we examined if artifacts in the data could be the reason for the low DEG recovery rate. Besides chromatin states with annotated function, chromatin state maps usually include a so–called background state that represents regions of no detectable signal (state number 18 labeled as “quiescent” in the CMM18 model). It is important to realize, though, that the interpretation of this background state is difficult. While it is conceivable that technical problems caused this lack of a signal in certain regions of the genome, it may be biologically meaningful in others. Moreover, the six canonical histone marks included in this study certainly cover a wide range of functionally important chromatin signals, but they do not represent the entire regulatory chromatin landscape. To give an example, the recently characterized H3K122ac histone modification is also found at active enhancers that lack the canonical H3K27ac marking [29]. Given these uncertainties, we opted for a conservative approach and considered the background state as not differential relative to all other chromatin states (see Methods and Figure 1).

We evaluated how many DEGs might not be recoverable under these conditions for the monocyte to macrophage comparison. For each of the 1110 DEGs that could not be recovered, we computed the percentage of the gene body length covered with the background state (averaged over all replicates in the respective groups). We found that close to a hundred genes that are covered to at least 60% with the background state are shared between the monocyte and the macrophage group (Figure 10A). At a higher threshold of 80% body coverage, this number drops to 35 genes. Given that this considers genes that are in the same uninformative chromatin state to roughly the same extent in all samples — and being differentially expressed at the same time — it seems justifiable to assume that the non–detection of these genes is not a limitation of SCIDDO. When focusing on the genes that are covered with the background state in either monocytes or macrophages, the numbers rise considerably (Figure 10B). 164 genes are above the lower threshold of at least 60% coverage, and when raising the threshold to at least 80% coverage, 72 genes are still affected. In this scenario, the non–detection of the DEGs is hence largely driven by the lack of a signal in one of the two sample groups.

**Figure 10:**
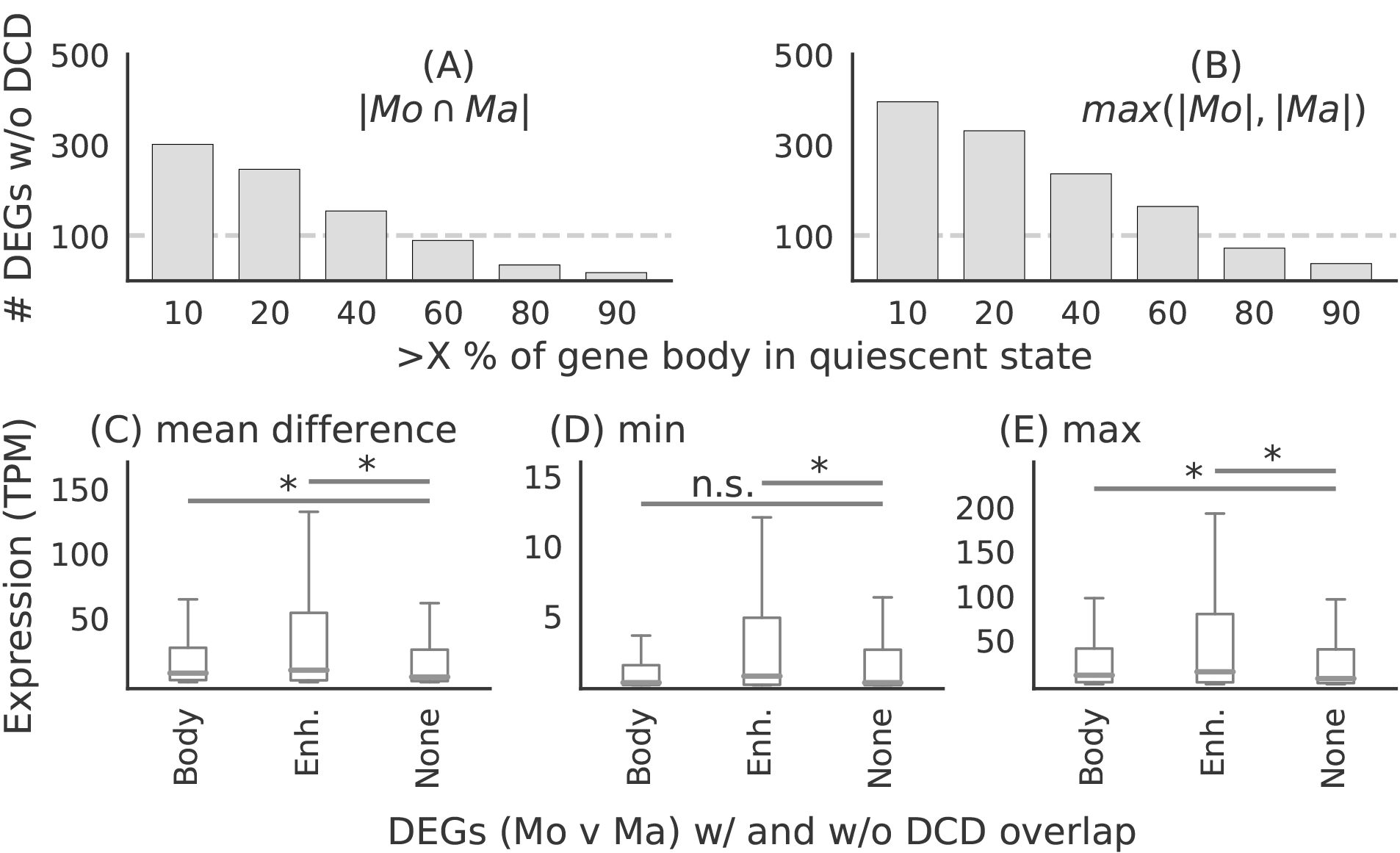
Uninformative chromatin state in gene bodies and moderate changes in expression complicate DEG recovery. Top: DEGs were binned according to the fraction of gene body covered with the background “quiescent” chromatin state (x–axis). (A) Height of bars depicts number of genes in intersection between monocyte and macrophage samples. (B) Height of bars depicts maximal number of genes either from monocyte or from macrophage samples. Bottom: DEGs were stratified according to DCD overlap in gene body/promoter (Body), or in at least one enhancer (Enh.) or no DCD overlap (None). Boxplots show distribution of gene expression values for absolute mean differences (C) between monocyte and macrophage samples, and for minimal expression (D) and for maximal expression (E) in any sample. Differences in magnitude were assessed using a two–sided Mann–Whitney–U test and considered significant (*) at *p* < 0.01 and not significant (n.s.) otherwise.

Considerations involving the background state might explain a few hundred cases of DEGs that could not be recovered by SCIDDO. It follows that a considerable amount of genes were assigned biologically meaningful chromatin states and yet were not detectable.

We hypothesized that a plausible cause for this could be a comparatively weak change in gene expression for non–detectable genes. When a gene is switched from “off” to “on”, a substantial change in the histone marking can be expected. However, if the gene is already actively transcribed and then simply upregulated, e.g., by activating additional enhancer elements (see Figures 6; Additional file 1: Figures S3 and S4), it is not obvious why this change in expression should lead to differential chromatin marking in the gene body. We tested this hypothesis by plotting the mean difference in expression, plus the minimal and maximal expression level in any sample, for all DEGs in the monocyte to macrophage comparison (Figure 10C–E). We split the genes into three groups based on DCD overlap in their gene body, in any associated enhancers but not in the body and no DCD overlap at all, i.e., the non–detectable genes. The mean change in gene expression is significantly higher in genes overlapping with a DCD compared to those genes that have no differential chromatin marking. Interestingly, the minimal expression level (Figure 10D) is still relatively high for those genes that show differential chromatin marking only in their enhancers. When relating the minimal to the maximal expression level (Figure 10D/E), the change in expression can be characterized as follows: genes with a DCD in their gene body jump from a low to a high expression level; genes with no DCD in their body but in their enhancer(s) show increased expression relative to an already high level, and genes with no DCD at all remain at a low to mildly elevated expression level. It should be pointed out that the implied directionality is supported by the observed expression changes for the monocyte to macrophage comparison (see Additional file 1: Figure S3).

There is a multitude of mechanisms beyond the chromatin level that can fine–tune gene expression [30–32]. Given that the DEGs lacking any sign of differential chromatin marking show also limited dynamics in their expression changes, we wondered whether there was any evidence of post–transcriptional control of these genes. As control group, we selected all genes that were not classified as differentially expressed but nevertheless showed signs of differential chromatin marking in their gene body (N=760 for the monocyte to macrophage comparison). We then plotted the number of annotated micro RNA targets using the TargetScan v7.2 [33] annotation for both groups of genes (Figure S6, bottom panel). There is indeed a small but statistically significant difference in the number of annotated micro RNA targets per gene between the two groups. This difference seems not to be caused by a difference in 3’–UTR length, where it is actually the group of DEGs without an overlapping DCD that has the larger 3’–UTR regions on average (Figure S6, top panel).

### SCIDDO affords direct interrogation of chromatin dynamics

A noteworthy feature of SCIDDO is the possibility to filter DCDs by chromatin dynamics. Given that chromatin states generated by the CMM18 model have been assigned meaningful labels (Additional file 1: Figure S1), users can exploit this easily interpretable annotation to filter DCDs. We used this feature in combination with external validation data to investigate if it is possible to identify enhancers that switch from an “on” to an “off” state between two cell types. To this end, we selected two sets of chromatin state labels as representing active and inactive enhancer states (see Methods). SCIDDO then uses these state labels to filter the DCDs and, by default, returns those subregions of a DCD where the chromatin change of interest can be observed between the selected cell types. It should be emphasized that, while the chromatin dynamics filtering is based on the identified DCDs, the individual subregions returned by SCIDDO cannot be statistically evaluated by computing an E–value. Subregions of a DCD can be as short as one or two genomic bins and, thus, the computed E–value of a subregion is unlikely to indicate statistical significance. For comparison, we downloaded several ENCODE peak datasets of the transcriptional co–activator EP300 (p300) for the cell line HepG2 (see Methods). Though EP300 is known to be highly predictive of tissue–specific enhancer activity [34], it cannot be assumed that all downloaded EP300 peaks mark active enhancers that are unique to HepG2, and are hence inactive in any other cell type. As a consequence, an exhaustive overlap between EP300 peaks and (switching) enhancer regions in DCDs cannot be expected. Instead, we hypothesized that it is more realistic to assume that any biologically meaningful enhancer switch within a DCD subregion should likely also show a change in EP300 occupancy. We investigated this hypothesis by plotting the count of EP300 peaks and their signal strength for all peaks generally overlapping DCDs, and for all peaks overlapping with DCD subregions showing enhancer switches from “on” to “off” and vice versa from “off” to “on” for the comparison of HepG2 to monocytes (Figure 11). There is a prominent difference both in absolute number of peaks and in signal strength for the two directions of enhancer switching. This example illustrates that SCIDDO can also offer support in downstream analysis by quickly identifying regions of specific and directed changes on the chromatin level.

**Figure 11:**
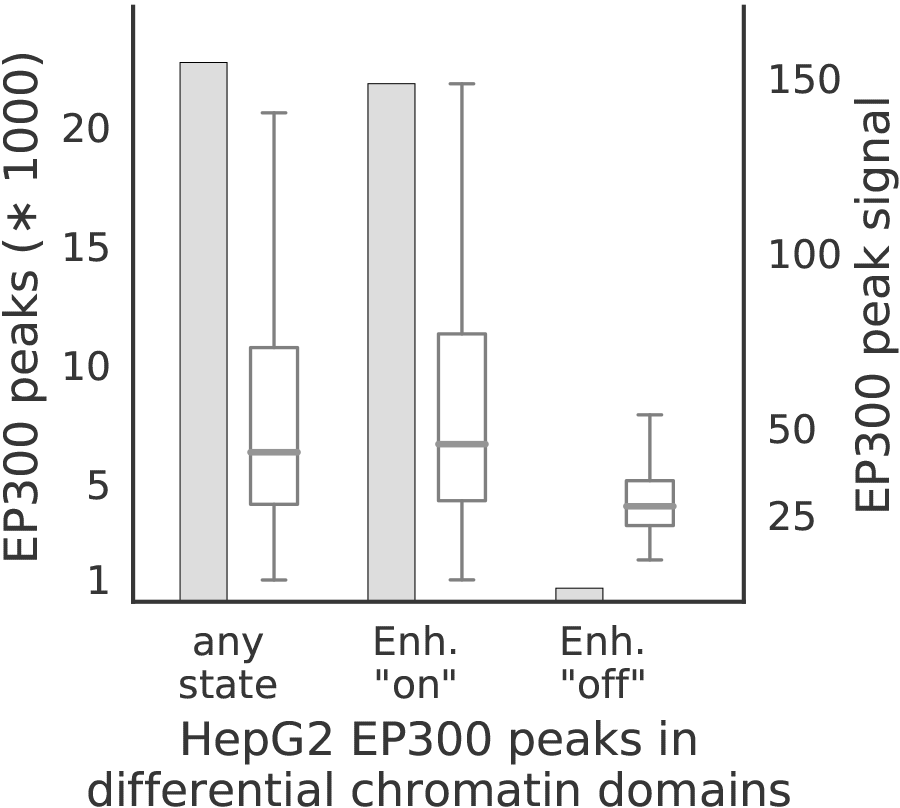
Chromatin dynamics at HepG2 enhancer elements. Height of the bars depicts total number of peaks overlapping DCDs (left y–axis) and box plots show distribution of the signal of the overlapping EP300 peaks (right y–axis). The three groups represent EP300 peaks overlapping with DCDs in general (left); with DCDs restricted to genomic locations showing an enhancer “on” state in HepG2 (middle); with DCDs restricted to genomic locations showing an enhancer “off” state in HepG2. For all three groups, the DCDs identified in the HepG2 to monocyte comparison were used.

### Differential chromatin domains recover differentially expressed genes with increased stability compared to individual histone marks

The number of available tools that use chromatin state maps as input for a differential analysis is limited. ChromDet [17] is designed for group comparisons with at least 5 to 10 replicates each (*personal communication*), and thus did not give results on our dataset. Similarly, ChromDiff [18] could not identify any differential chromatin marking (in genes), presumably due to lacking statistical power given the limited number of replicates in our dataset. The Chromswitch package [19] can only process one chromatin state at a time, which complicates direct and fair comparisons with the DCDs identified by SCIDDO.

We thus decided to compare SCIDDO to PePr [35], an established tool for the differential analysis of individual histone marks that can process replicated samples. This strategy has the advantage of reflecting the canonical “rule–based” approach of interpreting histone marks in well–characterized regulatory contexts, e.g., by determining enhancer activity based on the presence of H3K27ac peaks [36]. Specifically, we used PePr to perform a differential analysis for the same six sample group comparisons and evaluated PePr’s and SCIDDO’s performance for the task of detecting DEGs based on differential chromatin marking. To this end, we considered two different scenarios: first, genes overlapping with at least one differential chromatin domain (SCIDDO) or having at least one H3K36me3 peak in one cell type but none in the other cell type (PePr) were labeled as differentially expressed. This strategy could be applied to all 20,091 genes in our gene annotation (gene set G1). In the second scenario, differential chromatin in gene bodies was taken into account in the same way, but as an additional requirement, at least three annotated enhancers of a gene had to show differential chromatin marking (H3K27ac peaks for PePr) to label the gene as differentially expressed.

This reduced the number of genes in the evaluation set to 17,735 (88.3%; gene set G2), i.e., all genes that had at least three enhancers annotated. We compared the chromatin–based labeling of genes in sets G1 and G2 with the ground truth computed with DESeq2 [28]. While we settled for a fix threshold on gene expression fold change (> 2) and p–value (< 0.01) to identify DEGs throughout this study, we varied these values for the comparison between SCIDDO and PePr to examine the stability of their performance for different levels of differential expression stringency. We calculated accuracy and F1 score for all sample comparisons and the gene expression fold changes 0.5, 1, 2 and 4 and p–values 0.1, 0.05, 0.01 and 0.001 for the two gene sets G1 and G2 (Figure 12; Additional file 1: Figure S7). In summary, SCIDDO’s performance is superior to PePr. Averaged over all comparisons, SCIDDO shows an accuracy of 64.6% (G1) and 69.2% (G2) and a F1 score of 57.5% (G1) and 59.1% (G2) for the two different strategies of labeling a gene as differentially expressed. For PePr, the average performance scores are 57.6% (G1) and 57.7% (G2) accuracy and 54.6% (G1) and 54.7% (G2) F1 score.

**Figure 12:**
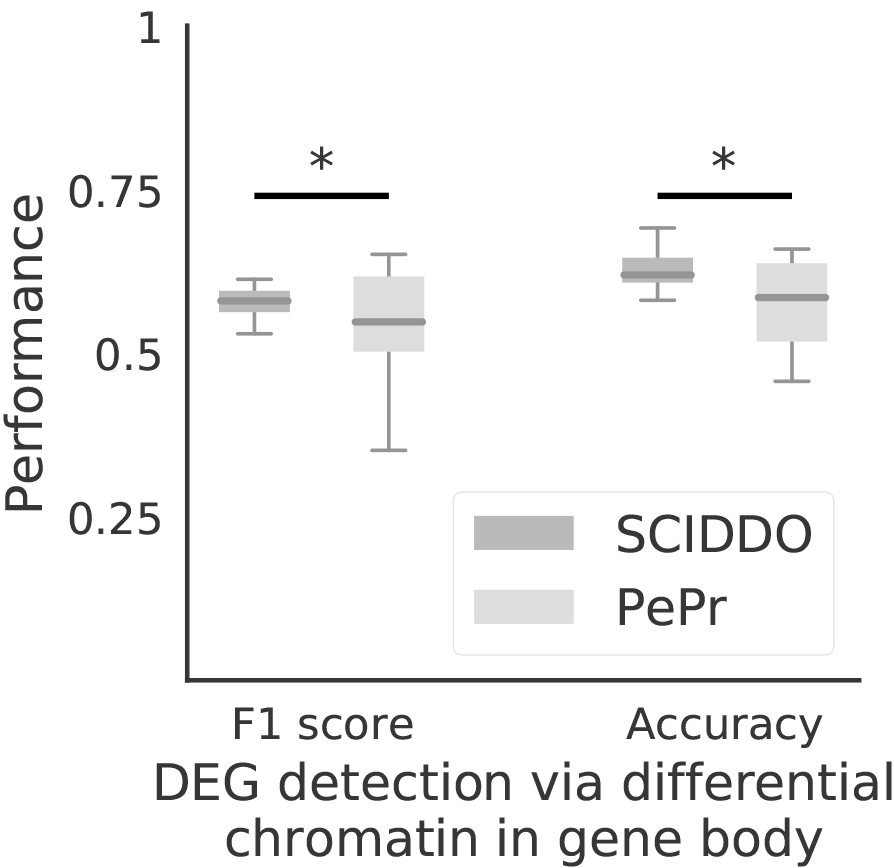
SCIDDO shows more stable performance at detecting DEGs. Box plots depict SCIDDO’s and PePr’s (light grey) performance of detecting DEGs quantified as F1 score (left) and as accuracy (right). Performance values are summarized over all sample group comparisons and for different thresholds on gene expression fold change (0.5, 1, 2 and 4) and on adjusted p–values (0.1, 0.05, 0.01 and 0.001) computed with DESeq2 to call DEGs. At least one DCD/differential H3K36me3 peak (PePr) was required in the gene body of a DEG to be considered detected on the chromatin level. Differences in performance were assessed with a one–sided Mann–Whitney–U test and considered significant “*” at *p* < 0.01.

## Discussion

The use of chromatin state segmentation maps for large–scale annotation and interpretation of reference epigenomes is well established in the field of computational epigenomics (see, e.g., [12, 37]). Nevertheless, comparatively little effort has been invested in the development of generally applicable software that assists researchers in exploiting these resources. To fill that gap, we developed SCIDDO, a new tool that implements a score–based approach for the fast detection of differential chromatin domains between potentially small groups of replicated samples.

The results presented above indicate that SCIDDO’s score–based approach is able to robustly identify consistent sets of differential chromatin candidate regions across individual biological replicate comparisons. This observation suggests that SCIDDO is well–equipped for the commonly encountered situation of limited replicate availability while still offering a statistically sound evaluation of the detected DCDs. Though the statistics implemented in SCIDDO do not afford a theory–driven evaluation of the detected DCDs, e.g., no suitable E–value threshold is motivated by the theory, we could validate our findings in several biologically meaningful ways. The considerable overlap between the detected DCDs and various regulatory annotation datasets (Figure 4) suggests a functional role for the identified DCDs that is in line with published studies [27, 38, 39]. By relating gene expression fold changes to DCD formation in gene bodies and gene–associated enhancers, we could show that this presumed functional role seems indeed to have a measurable effect on gene expression behavior (Figure 6, Additional file 1: Figures S3 and S4). Our findings conform to the established view that extensive chromatin changes in gene bodies as well as in gene–associated enhancers are good indicators of the expected gene expression fold change [40–42]. It should be emphasized that SCIDDO realizes this view on the interplay between chromatin changes and altered gene expression without directly quantifying differences on, e.g., the read count level. Nevertheless, SCIDDO is able to detect most DEGs (Figure 7) and shows a performance in such tasks that is on average superior and more stable compared to competing approaches which implement much more time–intensive strategies to differential chromatin analysis (Figure 12). Taken together, the evidence supports the conclusion that SCIDDO’s score–based approach to differential chromatin analysis discovers biologically meaningful and interpretable DCDs.

An observable trend in the dataset we analyzed is the more limited variation on the chromatin level with increasing cellular relatedness, e.g., what we have detailed for the monocyte to macrophage comparison. While this inverse relationship is plausible, it implies that there is a natural limit in “resolution” of differential chromatin state analyses that governs SCIDDO’s applicability in discerning cellular phenotypes or characterizing differentiation pathways. Although we did not investigate these potential limitations in depth, we collected multiple lines of evidence that illustrate various ways of how gene expression changes, and thus different cellular phenotypes, could be realized without necessarily leaving a detectable trace on the chromatin level (Figure 10, Additional file 1: Figure S6). One of these blind spots in chromatin state maps is the “quiescent” background state, i.e., the chromatin state without any detectable signal. If possible, a more fine–grained characterization of the background state would be a promising way of extending score–based differential chromatin analyses to cover even more regions of the (epi–) genome. To give an example, a widespread background state in gene bodies in only one sample group might be interpreted as biologically meaningful (cf. Figure 10B), and thus, an adapted scoring for the background state in this context could plausibly increase DEG recovery rates via DCD overlap.

Adaptations to the pairwise chromatin state scoring could be realized in a multitude of ways in future studies. While our approach based on the Jensen–Shannon–Divergence has the benefit of not being affected by biases in our dataset, which might be an issue for data–derived scoring systems, it is also not customized to any particular notion of differential chromatin. It is one of SCIDDO’s distinguishing features that the user can specify any scoring scheme that fulfills the statistical assumptions and use for differential chromatin analysis. It is thus conceivable to study only a specific repertoire of dynamic chromatin changes given an appropriately chosen scoring matrix, e.g., focusing on enhancers and ignoring transcribed regions. Apart from such specific objectives, it would also be intriguing to investigate if, for a given state segmentation model, generally applicable scoring systems could be derived that are sensitive to the degree of cellular relatedness. In analogy to genome sequence analysis [43], this could provide a different view on the dynamic epigenome in the course of cellular development.

## Conclusions

We developed SCIDDO as a versatile and fast tool to better exploit the abstract information stored in chromatin state segmentation maps. We presented evidence highlighting how the differential analysis of chromatin state maps can be conveniently used to identify differentially expressed genes or to characterize chromatin dynamics, and SCIDDO thus complements the bioinformatics tool box in exploratory epigenome studies. SCIDDO’s score–based approach lends itself to devising tailor–made scoring systems for specific analysis tasks and suggests a new way of interrogating chromatin data from a high–level perspective.

## Methods

### Experimental data overview

All analyses were carried out using the official IHEC human hg38/GRCh38 assembly. We selected the following high quality DEEP samples to include both closely related as well as more distantly related cell types in our analysis: two replicates of HepG2 (HG 1 and 2; Additional file 1: Table S1), two replicates of hepatocytes (He 2 and 3; Additional file 1: Table S1), three replicates of monocytes (Mo 1, 3, and 5 [20]) and two replicates of macrophages (Ma 3 and 5 [20]). All primary cell types were isolated from healthy, adult donors. For each replicate, we downloaded the DEEP reference alignments for six histone marks (H3K4me1, H3K4me3, H3K27ac, H3K27me3, H3K36me3, H3K9me3) and the corresponding Input control as BAM files (Additional file 1: Table S1). Additionally, we downloaded DEEP mRNA expression data for all samples as raw read FASTQ files (Additional file 1: Table S2). The hg38 genome reference was restricted to fully assembled auto– and gonosomes for all data preprocessing steps. The differential analysis with SCIDDO was then limited to autosomes and chromosome X to alleviate any effects arising from the uneven distribution of sexes in our dataset. Annotation data were likewise limited to the same set of chromosomes. The GeneHancer [44] enhancer annotation was licensed for academic use on 2017–05–30. The Gene–Hancer annotation was reduced to gene–enhancer pairs that could be mapped to gene identifiers in the GENCODE v21 annotation [45].

### Generation of chromatin state maps

Following IHEC recommendations, all histone BAM files were filtered using Sambamba v0.6.6 [46] to exclude low quality reads (mapping quality ≥ 5; no duplicated, unmapped or non–primary reads/alignments). These filtered BAM files were used as input to generate chromatin state segmentation maps for all samples. We used the pre–trained 18–state ChromHMM (CMM18) model provided by the Roadmap Epigenomics Mapping Consortium (REMC [21]). We decided to use this pre–trained model because it has been carefully designed using the large compendium of epigenomes generated by the REMC. We thus assumed that this model robustly captures chromatin states irrespective of the biological source of the samples at hand. As an additional benefit, the chromatin states of the CMM18 model were functionally characterized and labeled by the REMC to make interpretation of the state segmentation maps straightforward (Additional file 1: Figure S1, Table S3). We executed version 1.12 of ChromHMM with commands `BinarizeBam -b 200` and `MakeSegmentation -b 200` and otherwise default parameters to create the state segmentation maps.

### Differential gene expression analysis

Gene expression estimates per replicate were computed with Salmon v0.9.1 [47] using the GENCODE v21 [45] annotation for protein coding genes. For each gene in the GENCODE reference, we extracted genomic coordinates for the gene body (5’ to 3’ end) and for the promoter (−2500 bp to +500 bp around the 5’ end) using custom scripts. After expression quantification, we used DESeq2 v1.18.1 [28, 48] to obtain differential expression estimates for all six possible pairs of sample replicate groups in our dataset. We split the DESeq2 results into groups of differentially expressed genes (DEGs) and non–differentially expressed genes (stable genes) based on an absolute log2 fold change in expression of at least 2 and a multiple testing corrected p–value of less than 0.01.

### Differential histone peak calling

We selected PePr [35] as a current state–of–the–art tool for differential chromatin analysis as a reference to compare to. We executed PePr v1.1.18 to perform differential analysis including postprocessing for all six possible pairs of sample replicate groups in our dataset. All available replicates were processed in a single run of PePr for each comparison. PePr was executed with histone peak type set to `broad` for the mark H3K36me3, and otherwise default parameters. The resulting histone peak sets were filtered to peaks with a q–value of less than 0.01 using custom scripts.

### Chromatin dynamics at EP300 peaks

EP300 peak datasets for HepG2 were downloaded from ENCODE [3] (ENCFF674QCU and ENCFF806JJS) and merged using bedtools v2.26.0 [49]. For the chromatin dynamics filtering, chromatin states 7–11 (genic, active and weak enhancers) were considered as enhancer “on” states, and chromatin states 13, and 15–17 (heterochromatin, bivalent enhancer and polycomb repression) were considered as enhancer “off” states.

### Statistical background for SCIDDO

The theory behind the statistical evaluation available in SCIDDO has been developed in the context of biological sequence analysis, e.g., to identify runs of hydrophobic amino acids in protein sequences [50, 51]. Since the theory was left unaltered, we give only a compact overview to introduce the necessary concepts and nomenclature. The chromatin state maps of each sample in the SCIDDO dataset can be represented as a sequence *X* = {*x*_1_… *x*_*p*_… *x*_*n*_}. Here, the *x*_*p*_ are assumed to be i.i.d. random variables over an alphabet *A* and *n* is the length of the sequence. In our case, |*A*| = 18 representing the 18 different chromatin states of the CMM18 model. Each pair of states (*a*^*i*^, *a*^*j*^) is assigned a score *s*^*ij*^ where *s*^*ij*^ < 0 indicates state similarity (uninteresting regions) and *s*^*ij*^ > 0 indicates state dissimilarity (interesting regions; see below for derivation of the *s*^*ij*^). We omit the superscript *ij* in the following to improve readability. When comparing two chromatin state maps *X*, *Y*, each state pairing (*x*_*p*_, *y*_*p*_) is assigned the respective score *s* as defined above. This results in a sequence of scores *S* = {*s*_1_ … *s*_*n*_} that is scanned for subsegments of highest cumulative score. This approach is called local score computation and can be done efficiently with a linear time algorithm [52]. The set of all maximal scoring disjoint segments returned by this algorithm represents the set of candidate regions for the respective chromatin state map comparison. The (unnormalized) raw score *R* of a candidate region is simply defined as the sum over all scores in the candidate region 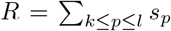 where *k* and *l* indicate the position of the leftmost and of the rightmost genomic bin included in the candidate region. These cumulative scores have to be normalized to account for the fact that higher scores have a higher chance of occurring with increasing sequence length. This normalization step requires the estimation of two statistical parameters λ and *K* (for detailed derivation of these parameters, see [51]). Since both λ and *K* lack a biologically meaningful interpretation, they can be simply thought of as scaling parameters for the scoring system and the search space. For this parameter estimation, SCIDDO relies on the routines implemented in BLAST v2.7.1 [53]. Additionally, four assumptions are needed for the theory to be applicable, which then allows to model the limiting behavior of the score distribution as Gumbel–type extreme value distribution (see [51], Figure 2):

1. The sequences are infinitely long
2. The *x*_*p*_ are i.i.d. random variables
3. A positive score must be possible
4. The expected score is negative

Assumptions 1. and 2. of course do not apply to any biological sequence, but are needed for reasons of mathematical tractability [51]. Assumptions 3. and 4. are tested by SCIDDO before starting the actual analysis, safeguarding against errors in the statistical evaluation. Under these conditions, the Expect value (E) for a DCD with raw score *R* is then calculated as

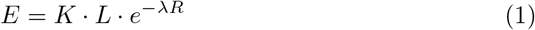

where the factor *L* is the length of the chromosomal sequence adapted for replicate–variation. Since SCIDDO has been designed to compare (small) groups of replicates against each other, we adapted the calculation of the total length of the sequence. Intuitively, adding more and more biological replicates to a group of samples does not linearly increase the amount of information contained in the respective group. At some point, all biologically meaningful variation has been sampled and, ignoring technical artifacts and stochastic effects, no new chromatin states should be observed at any position of the genome. Based on this consideration, for each additional replicate in a group of samples, SCIDDO only adds those positions to the total sequence length that show a new chromatin state compared to all other biological replicates already in the group. A complete description of this computation is given as pseudocode in Additional file 1: Algorithm 1.

### Fit of random scores to Gumbel–type extreme value distribution

The calculation of the E–value as described above assumes a null model of random sequences. Following the theory (cf. Theorem 1 in [51] and examples in [22]) the normalized maximal scores should follow a Gumbel–type extreme value distribution when comparing random state sequences, in the limit of the sequence length *n*. Since SCIDDO supports the use of customized scoring schemes, it also supports the user in assessing if the chosen scoring scheme follows this theoretical assumption. To that end, SCIDDO scans the randomly shuffled chromatin state maps of all sample pairs for high scoring subsegments and retains only the maximally scoring subsegment per chromosome; if several segments with identical scores emerge, only the first one is kept. This process is iterated until a pre-specified number of these “random” scores have been found. The scores underlying Figure 2A have been generated in that way. The user can then use these “random” scores and, e.g., assess their fit to a Gumbel–type extreme value distribution following our example in Figure 2A. Notably, in Figure 2A, we jointly fitted all “random” scores of all chromosomes to simplify the visualization.

### Derivation of pairwise chromatin state similarity scores

The theoretical considerations presented in the previous section do not enforce the use of complicated scoring systems that are in turn well–grounded in theory, e.g., rather simple “match/mismatch” or empirically derived scoring systems can be used if considered appropriate [50]. We thus decided to use the emission probability vectors of the 18 chromatin states (= the hidden states of the ChromHMM Hidden Markov Model) to compute pairwise similarity scores. The state emissions 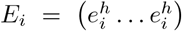 for state *a*_*i*_ represent a probability distribution over the observed outputs, i.e., over the observed six histone modifications *h*. This motivated using the symmetric Jensen–Shannon–Divergence (JSD) [54] to compute chromatin state similarities

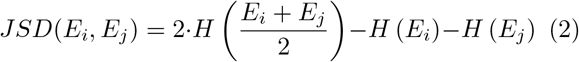

where *H* is the Shannon entropy

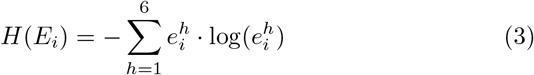

Since the JSD has a lower bound of 0, the pairwise similarities for each state were shifted by subtracting the mean JSD. This resulted in negative scores for similar states (JSD near zero) and positive scores for dissimilar states. Scores are commonly represented by integer values, which we realized by multiplying the real–valued scores by a factor of 10 and rounding them to integers afterwards. As mentioned above, SCIDDO checks the adherence to assumptions 3. and 4. for any custom scoring scheme such as our JSD–derived one to ensure applicability of the Karlin–Altschul statistics before starting a differential analysis.

A peculiarity of chromatin state maps is the so–called background state (state 18 labeled as “quiescent” in the CMM18 model). This state represents the lack of any detectable signal in the input data. As it is *a priori* impossible to identify the true source for this lack of a signal, i.e., it could be a technical artifact or biologically meaningful, the background state needs to be handled with special care in the interpretation of chromatin state maps. We decided to implement a cautious strategy and replaced all pairwise state similarities involving the background state with the minimal score generated with our JSD–based approach. In other words, the background state is similar, i.e., not differential relative to all other chromatin states. We opted for this strategy to avoid finding differential chromatin domains that are dominated by the background state and could thus be challenging to interpret.

## List of Abbreviations

bp: base pair(s)
DCD: differential chromatin domain
DCS: differential chromatin score
tf: transcription factor
DEG: differentially expressed gene

## Declarations

### Availability of data and material

The DEEP sequencing datasets analyzed in this study are not publicly available due to patient privacy. Access to the raw data can be requested under www.ebi.ac.uk/ega/dacs/EGAC00001000179.

Pipeline code to reproduce all results and figures of this study is available under doi.org/10.17617/1.6K. The source code of the SCIDDO tool is available under github.com/ptrebert/sciddo.

### Author’s contributions

P.E. and M.H.S. conceptualized the project; P.E. carried out the implementation work, analyzed the data and wrote the first draft of the manuscript; all authors contributed to the writing of the manuscript. All authors read and approved the final manuscript.

## Acknowledgements

We acknowledge the DEEP consortium for providing the epigenome and transcriptome data. We thank Thomas Lengauer for critical reading of the manuscript. Additionally, we thank Prabhav Kalaghatgi, Nora K. Speicher and Martin Vingron for helpful discussions.

## Additional Files

Additional file 1 — Supplementary material PDF format

Contains all supplementary figures and tables.

